# Onset–Rime Orthographic Representations in Early Literacy Development: A Neural Link Between Spelling and Reading

**DOI:** 10.64898/2026.10.01.756116

**Authors:** Fang Wang, Lindsey R. Hasak, Vani Dewan, Bruce D. McCandliss

**Affiliations:** Department of Linguistics and Translation, City University of Hong Kong, Kowloon, Hong Kong; Graduate School of Education, Stanford University, Stanford, CA, USA; Department of Psychology, University of South California, CA, USA; Wu Tsai Neurosciences Institute, Stanford, CA, USA

**Author notes:** Corresponding author: Bruce McCandliss,. Graduate School of Education, 485 Lasuen Mall, Stanford, CA 94305 USA.

**Keywords:** Orthographic Knowledge Onset–Rime Structure Spelling and Reading Neural Sensitivity Fast Periodic Visual Stimulation

## Abstract

Visual orthographic knowledge plays a critical role in early literacy development and may provide a shared foundation for reading and spelling, yet the nature of its underlying representations remains debated. Behavioral measures offer indirect evidence and may be influenced by other metalinguistic processes. It remains unclear whether beginning readers encode larger orthographic structures, such as onset–rime units, beyond individual letters and smaller letter combinations but below the whole-word level. It is also unknown whether neural sensitivity to these structures helps explain the relationship between spelling and reading. To address these questions, we used a fast periodic visual stimulation (FPVS) paradigm in first- and second-grade children to contrast pseudowords constructed with the typical ordering of onset followed by rime elements (i.e., onset–rime) against matched strings in which the same elements appeared in the reverse order (i.e., rime–onset). Neural sensitivity to on-set–rime orthographic structure was associated with both spelling ability and word-reading efficiency and, critically, statistically mediated the relationship between these two literacy skills. These relationships remained significant after controlling for age and removing influential observations, and sensitivity to onset–rime orthographic structure explained additional variance in word-reading efficiency beyond spelling ability and age. Together, these findings provide neurophysiological evidence for onset–rime structure as an important sublexical orthographic unit in beginning readers, highlighting its potential role in linking spelling with efficient word recognition during early reading development.

## 1 Introduction

Reading and spelling are traditionally considered as “two sides of the same coin” in literacy development, as they are highly correlated and follow similar developmental trajectories in early readers (Ehri, 2000; Perfetti, 1997). Behavioral studies have consistently shown that spelling ability is strongly associated with word reading and decoding fluency (Graham & Santangelo, 2014; Treiman, 2017). Moreover, longitudinal and intervention studies suggest that spelling may contribute uniquely to reading development above and beyond phonological awareness, letter-sound knowledge, and other cognitive–linguistic abilities (Ouellette & Śeńechal, 2017; Śeńechal, 2019; Treiman et al., 2019).

One explanation for the close relationship between reading and spelling is that both skills rely on a shared system of orthographic knowledge (Chang et al., 2018; Ehri, 2000; Mather & Jaffe, 2021). Orthographic knowledge refers to knowledge of how spoken language is represented in written form, including the visual patterns, letter combinations, and structural regularities that characterize a writing system (Apel, 2011; Deacon et al., 2012). Such knowledge enables readers and spellers to recognize, produce, and generalize written forms across orthographically related words (Conrad et al., 2013; Treiman, 2013; Zaric et al., 2021). Accordingly, orthographic knowledge is widely considered as a core foundation of literacy development across writing systems and has been proposed to contribute to variations in reading and spelling abilities (Ehri, 2000; Perfetti, 2007; McBride et al., 2022; McBride, 2025).

Although decades of behavioral research have established orthographic knowledge as a common foundation linking reading and spelling (Bosse, 2015; Ehri, 2014), remarkably little is known about how this knowledge is represented in the developing brain. Existing evidence for the underlying orthographic representations has come primarily from behavioral measures, making it difficult to characterize their underlying organization directly. Specifically, existing behavioral measures, such as orthographic choice, spelling judgments, and pseudoword reading, provide only indirect evidence about visual orthographic representations because they rely on overt decisions and responses (Mather & Jaffe, 2021; Apel, 2011; Deacon et al., 2012). As a result, performance on these tasks reflects not only orthographic processing itself but also other cognitive and linguistic abilities, including metalinguistic awareness (Treiman & Kessler, 2014), phonological and speech-production demands (Share, 1995; Shahar-Yames & Share, 2008), working memory (Berninger et al., 2006), and attention control (Thomson et al., 2005). Therefore, objective neural measures provide an opportunity to move beyond indirect behavioral inference by directly probing the neural representations that support reading and spelling.

Investigating visual orthographic representations at the neural level requires understanding how these representations are organized. Many influential models of visual word recognition, including interactive activation, dual-route cascaded (DRC), and connectionist models, have primarily focused on orthographic processing at the level of individual letters and whole-word representations (Coltheart et al., 2001; McClelland & Rumelhart, 1981; Seidenberg & McClelland, 1989). For example, the DRC model proposes that skilled reading can be supported by stored *whole-word* representations or serial *letter-by-letter* decoding (Coltheart et al., 2001). At the neural level, increasing evidence suggests that reading development is accompanied by the emergence of specialized visual responses to orthographic information within the ventral occipito-temporal system (Dehaene-Lambertz et al., 2018; McCandliss et al., 2003; Turkeltaub et al., 2003a). Neural sensitivity to orthographic structure has been consistently linked to reading skill and may provide an objective index of automatic orthographic processing (Araújo et al., 2015; Lochy et al., 2025; Lutz et al., 2024; Wang et al., 2024). These findings suggest that neural measures provide a unique opportunity to investigate the organization of visual orthographic representations directly. However, most of these neural studies have focused on local or lower-level aspects of orthographic processing, such as individual letters, short consonant–vowel–consonant (CVC) strings, orthographic legality, or on broader distinctions such as lexicality (Gigleux et al., 2026; Lochy et al., 2025; Wang et al., 2023, 2024).

This leaves open the possibility that the developing orthographic system may also encode *intermediate* orthographic representations that capture recurring patterns across words. Such intermediate representations could provide a bridge between individual letters and whole words by representing orthographic units larger than individual letters or simple letter combinations (e.g., bigrams and trigrams), yet smaller than whole-word representations (Dehaene et al., 2005; Grainger & Van Heuven, 2003). In turn, these intermediate representations may support efficient recognition of familiar words while enabling readers to generalize orthographic knowledge across unfamiliar words that share common orthographic patterns (Deacon & Bryant, 2006). This possibility is particularly important in English, where phoneme–grapheme correspondences are often inconsistent and do not follow simple one-to-one mappings (Frost et al., 2005; Share, 2008; Ziegler & Goswami, 2005). Under such conditions, sensitivity to recurring orthographic patterns may allow readers to make use of regularities that extend beyond individual grapheme–phoneme correspondences (Deacon & Bryant, 2006). However, the neural organization of such intermediate orthographic representations in developing readers remains poorly understood. It also remains unclear whether neural sensitivity to these representations may help explain the close relationship between reading and spelling.

One candidate for such an intermediate orthographic representation is onset–rime structure. Onset–rime structure has long been considered a psychologically meaningful unit in reading and spelling, particularly during early literacy development, and plays a central role in theories proposing that generalization across orthographically related word families contributes to the development of orthographic knowledge (Castles et al., 2018; Greaney et al., 1997; Treiman, 2020). Many English words can be grouped into word families that share a common rime (e.g., c*at*, h*at*, m*at*, s*at*), making onset–rime regularities a common feature of written English (Goswami, 1990, 1991). Behavioral studies further suggest that developing readers are sensitive to this organization: children can use shared rime patterns to generalize across words, and the learning of spelling patterns is facilitated when those patterns correspond to onset–rime structure (Goswami, 1991; Goswami & Mead, 1992; Greaney et al., 1997; Wise et al., 1990). Together, these findings suggest that onset–rime structure may be encoded as an intermediate orthographic representation that captures reusable patterns shared across words. However, most evidence for onset–rime structure has come from behavioral studies, and it remains unclear whether this structure is automatically encoded in children’s visual orthographic representations at the neural level and whether neural sensitivity to this structure may help explain the relationship between reading and spelling.

To directly test this possibility, we used fast periodic visual stimulation, an objective EEG paradigm for measuring automatic visual discrimination, to examine neural sensitivity to onset–rime structure in first- and second-grade children. (Rossion, 2014; Norcia et al., 2015). Specifically, we examined neural responses to strings constructed from onset (e.g., kn) and rime units (e.g., ight) derived from real English words (e.g., knight). Neural responses to strings with a conventional onset–rime structure (e.g., glatch, snight) were compared with matched strings containing the same orthographic units arranged in a rime–onset order (e.g., atchgl, ightsn). If onset–rime orthographic structure is encoded within visual orthographic representations, we expected stronger neural responses to conventionally organized onset–rime strings than to matched rime–onset strings. This contrast therefore provided a direct neural test of whether developing readers encode onset–rime structure as part of an intermediate orthographic representation. Critically, we examined whether neural sensitivity to onset–rime orthographic structure mediates the well-established relationship between spelling and reading, and may therefore help explain why these two literacy skills are closely related.

## 2 Methods

### 2.1 Ethics Statement

This research was approved by the Institutional Review Board of Stanford University. All participants delivered written informed consent prior to the study after the experimental protocol was explained.

### 2.2 Participants

Sixty-five English speaking first- and second-grade students (33 girls) at an independent school participated in this study. Of these, fifty-nine participated in the EEG session. Nine participants were excluded due to data quality (*N* = 7) and technical (*N* = 2) issues, resulting in a final sample of *N* = 50 (m = 7.46 years, sd = 0.71 years).

### 2.3 Behavioral Assessments

Forty-eight of the 50 participants who completed the EEG session also completed an approximately 30-minute behavioral assessment afterwards (summarized in Table 1). All participants were tested for handedness using the Edinburgh Handedness Inventory (Oldfield, 1971). Children also completed measures of phonological awareness and rapid naming, including the Rapid Automatized Naming (letters and colors) subtests from the Comprehensive Test of Phonological Processing, Second Edition (CTOPP-2; Wagner et al. (2013)). Spelling and decoding abilities were measured using the spelling and letter-word identification (LWID) subtests of Woodcock-Johnson Tests of Achievement, Fourth Edition (WJ-IV; Schrank et al. (2014)). Word reading fluency was assessed using the Test of Word Reading Efficiency, Second Edition (TOWRE-2; Torgesen et al. (2012)), while silent reading efficiency and reading comprehension fluency were assessed using the Test of Silent Reading Efficiency and Comprehension (TOSREC; Wagner et al. (2010)).

**Table 1:** Performance on behavioral assessments. . Values are range and mean(*±SD*). RAN: Time (ms) used to quickly and accurately name all stimuli (e.g., letters or colors) on a test form. WJ-Spelling: Number of correctly spelled words until six consecutive errors occur. WJ-LWID: Number of correctly named letters and words on a test form. TOWRE: Number of real words and pronounceable nonwords read in 45 seconds. TOSREC: Number of sentences read and correctly verified within three minutes. Note: All scores are raw scores.

|  |  |  |  |
| --- | --- | --- | --- |
|  | Sex (female/male) | 26/22 |  |
|  | Handedness (right/left/ambidextrous) | 38/3/7 |  |
|  | Age in years | 6-8 | 7.5(±0.7) |
|  | Rapid automatic naming of colors (RANcolor) | 21-61 | 35.9(±10.4) |
|  | Rapid automatic naming of letters (RANletter) | 13-49 | 24.4(±7.5) |
|  | Word spelling ability (WJ-Spelling) | 14-43 | 24.0(±8.8) |
|  | Word decoding ability (WJ-LWID) | 24-68 | 47.0(±16.3) |
|  | Test of word reading efficiency (TOWRE) | 19-134 | 106.1(±21.1) |
|  | Test of silent reading efficiency and comprehension (TOSREC) | 1-60 | 28.5(±16.5) |

### 2.4 Study Procedure

In collaboration with administrators responsible for the school’s literacy program and lower-grade teachers, 36 words were drawn from late second-grade and early third-grade curricula to ensure that they were both challenging and appropriate for the students. These words comprised nine word families (i.e., rime units; e.g., -erve, -udge, and -atch), with four items per family (i.e., orthographic neighbors sharing the same rime unit but different onsets; e.g., nerve, verve, serve, swerve).

We created well-matched pseudoword (PW; e.g., glight, snatch) and nonword (NW; e.g., atchgl, ightsn) stimuli by recombining onset (e.g., pl) and rime (e.g., ind) units derived from real English words list. PWs preserved the conventional onset–rime organization, whereas NWs consisted of the same orthographic units arranged in the reverse rime–onset order. EEG data were recorded while participants were presented with PW and NW stimuli in an alternating sequence at a base frequency of 3 Hz (one string every 333 ms, three stimuli per second) and an oddball frequency of 1 Hz (one oddball per second). This means that three stimuli per second were presented at a constant rate (3 Hz) and the three stimuli presented in a given second always comprised two NWs followed by one PW (1 Hz).

A total of 30 trials were presented, each lasting 12 seconds, resulting in 360 seconds of continuous stimulation per participant. The 30 trials were divided into three blocks, which were completed in a pseudorandomized order across participants to minimize potential order effects. Feedback and breaks were provided between trials and blocks as needed.

### 2.5 EEG Recording Procedure

Participants were seated approximately 1 m from a computer monitor in a dimly lit room. A brief practice session was conducted prior to the experiment to ensure familiarity with the task procedure. EEG data were collected using 128-sensor HydroCell arrays (MagstimEGI), Electrical Geodesics NetAmp300, and NetStation 5.4.2 software, while stimuli were presented via an in-house software. Data were recorded against Cz reference, sampled at 500 Hz. Impedances were kept below 50 kΩ.

To maintain participants’ attention throughout the experiment, a repetition detection task was used. Participants were instructed to press a button on a response pad with their dominant hand when a “target” stimulus appeared three times consecutively. To enhance engagement and align with the school-wide theme, the targets were sea-animal words: shrimp, otter, whale, and guppy. Feedback was provided after each trial using cartoon images corresponding to the target words, and participants were asked whether they had seen any of the images.

Each block’s stimulation sequence began with a blank screen with a jittered duration ranging from 1500 to 2500 ms. Each block comprised 10 trials (12 seconds per trial), resulting in 30 trials across all three blocks. Within each trial, stimuli were presented as one NW-NW-PW triplet per second, for a total of 12 triplets per trial. Of the 10 trials presented pseudo-randomly presented within each block, four were target trials (i.e., contained target stimuli): in two trials, the repeated stimuli appeared at the end of the trial (“terminal” trial), whereas in the other two trials, they occurred at random positions within the trial (“catch” trial). Each target trial contained only one target. The remaining six trials contained no repeated stimuli. Due to excessive movements associated with the button presses, the two catch trials were systematically removed from further data analyses, whereas the two terminal trials were still included because the movements happened at the end of the trial. In total, 8 trials per block (24 trials overall) were included in the final analyses.

Each EEG session took approximately 40 minutes, including setup and between-trial breaks. Each child received a small token gift (e.g. stickers) after the study as compensation.

### 2.6 EEG Preprocessing

Recorded EEG data were bandpass filtered offline (zero-phase filter, 0.3–50 Hz) using Net Station Waveform Tools. Data were then imported into the in-house signal processing software for further pre-processing.

During pre-processing, EEG data were resampled to 420 Hz to ensure an integer number of time samples per video frame at a 60 Hz frame rate, as well as an integer number of frames per cycle for the present stimulation frequencies (i.e., 3 Hz). Sensors were identified as noisy if more than 15% of their samples exceeded an amplitude threshold of 60 *µ*V; these sensors were interpolated using the average of the six nearest neighboring sensors. Continuous EEG data were subsequently re-referenced to average reference (Lehmann & Skrandies, 1980) and segmented into 1-s epochs. Artifact rejection was conducted on a sensor-by-sensor basis, excluding epochs in which more than 10% of samples exceeded 60 *µ*V or any individual sample exceeding the artifact threshold of 60 *µ*V (e.g., due to eye blinks, eye movements, or body movements). The first and last 1-second epochs of each trial—each comprising a NW-NW-PW triplet—were excluded from further analysis, to minimize onset-related transient responses and blinking artifacts at the beginning and end of each trial. EEG signals were then filtered in the time domain using Recursive Least Squares (RLS) filters (Tang & Norcia, 1995) optimized for each frequency of interest, and transformed into complex Fourier coefficients for subsequent frequency-domain analyses. Complex-valued RLS outputs were decomposed into real and imaginary coefficients for input to the spatial filtering computations of Reliable Component Analysis (RCA), as described below.

### 2.7 Analysis of Behavioral Data

Behavioral responses for the repetition detection task served to monitor participants’ attention during EEG recording. We computed d’ based on the z-transformed probabilities of hits and false alarms (Macmillan & Creelman, 2004)

### 2.8 Analysis of EEG Data

Sensor-space EEG data were spatially filtered using Reliable Components Analysis (RCA, Dmochowski et al. (2012, 2015)), a matrix decomposition technique that derives spatial filters maximizing trial-to-trial covariance relative to within-trial covariance, thereby isolating phase-locked neural responses across repeated stimulations. Given the stable response phases characteristic of SSVEPs, RCA decomposed the 128-sensor array into a small set of reliable components (RCs) reflecting maximally consistent neural activity. For each RCA computation, the first five reliable components were extracted from 1-s epochs of sensor-by-feature data, including real and imaginary Fourier coefficients at frequencies of interest. To examine neural responses to the contrast between onset–rime and rime–onset stimuli, RCA was computed across oddball frequency and its harmonics (1, 2, 4, 5, 7, and 8 Hz), excluding overlapping base harmonics (3, 6, and 9 Hz). To examine general visual processing, RCA was conducted using base frequency and its harmonics (3, 6, and 9 Hz).

To quantify overall response amplitude across harmonics, we calculated the root sum of squares (RSS; hereafter referred to as RSS amplitude) by taking the square root of the summed squared amplitudes across harmonics.

Response latency (i.e., cortical processing speed) estimates were obtained from the phase information of harmonics showing significant responses. Specifically, phase values were modeled as a function of harmonic frequency using linear fitting implemented with MATLAB’s polyfit function (Norcia et al., 2020). Neural response latency was then derived from the slope of the fitted phase–frequency relationship.

### 2.9 Analysis of Brain-Behavior Relationships

Brain–behavior analyses were conducted to examine individual variability in the relationship between RSS amplitude and literacy skills. Behavioral measures included raw scores on WJ Spelling and TOWRE, indexing spelling ability and word-reading efficiency, respectively. Linear regression analyses were performed to assess the associations between RSS amplitude and spelling and reading measures. Multiple comparisons were controlled using false discovery rate (FDR) correction. When significant correlations were observed, influential observations were identified using Cook’s Distance (Cook, 1977) and excluded if they exceeded the 4/n threshold, where n represents the total number of observations. Correlation analyses were then repeated to determine whether the observed relationships remained significant after the removal of influential data points. Because age is strongly associated with reading development (Nahavandi et al., 2020), multiple linear regression analyses were further conducted with age included as a covariate to assess whether RSS amplitude remained significantly associated with spelling and reading measures after controlling for age.

To further investigate the relationship between word spelling and word-reading efficiency, mediation analyses were conducted to test whether neural orthographic sensitivity mediated the association between these two literacy skills. This analysis was motivated by the established relationship between spelling and reading, as well as the observed associations between RSS amplitude and both behavioral measures. Indirect effects were examined using a nonparametric bootstrap procedure with 1,000 resamples to estimate confidence intervals and p-values. Standardized regression coefficients were calculated and reported as indices of effect size.

To determine whether whether neural sensitivity to onset–rime orthographic structure explained unique variance in word-reading efficiency beyond spelling ability and age, hierarchical linear regression analyses were conducted. In Model 1, TOWRE scores were predicted from WJ Spelling and age. In Model 2, neural orthographic sensitivity (RSS amplitude) was included as an additional predictor. Model improvement was evaluated using nested model comparisons and the significance of regression coefficients in the full model.

## 3 Results

### 3.1 Robust neural sensitivity to onset–rime structure in early readers

Robust neural responses to orthographic structure were observed, as indexed by significant onset–rime vs. rime–onset responses across harmonics over left occipito-temporal regions (Figure 2). Figure 2A represents neural responses to onset–rime structure. Topographies (Figure 2A.1) and bar plots (Figure 2A.2) illustrate significant neural responses over left occipito-temporal regions to orthographic structure at all oddball harmonics except for 7 Hz. Phase (*π*) by harmonic frequency functions were used to estimate response latency in milliseconds (ms), yielding an estimated latency of 194.08*±*9.7 ms (Figure 2A.3), which is consistent with previous literature showing slightly delayed N170 latencies in children (*∼*190–200 ms). To test the stability of neural responses to onset–rime structure, we examined split-half reliability across stimulus sets. Figure 2A.4 demonstrates high split-half reliability, indicating that these responses were highly stable across stimulus sets.

**Figure 1:**
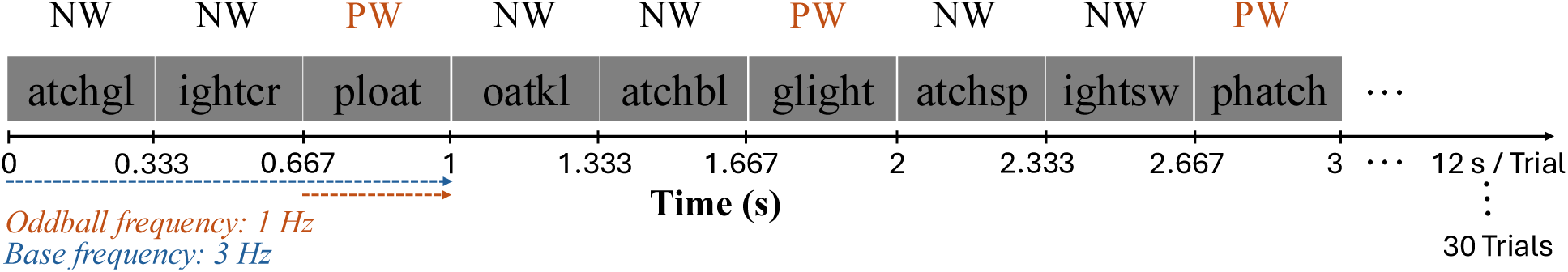
Examples of stimuli presented in the experiment. 1 Hz onset–rime pseudoword (PW) oddballs were embedded in a 3 Hz stream of rime–onset nonwords (NW). Each trial consisted of 12 seconds of stimulation (three stimuli presented per second, with two NWs followed by one PW), with 30 trials presented in total. The trials were divided into three blocks, which were presented in a pseudorandomized order across participants. Feedback and breaks were provided between trials and blocks as needed.

**Figure 2:**
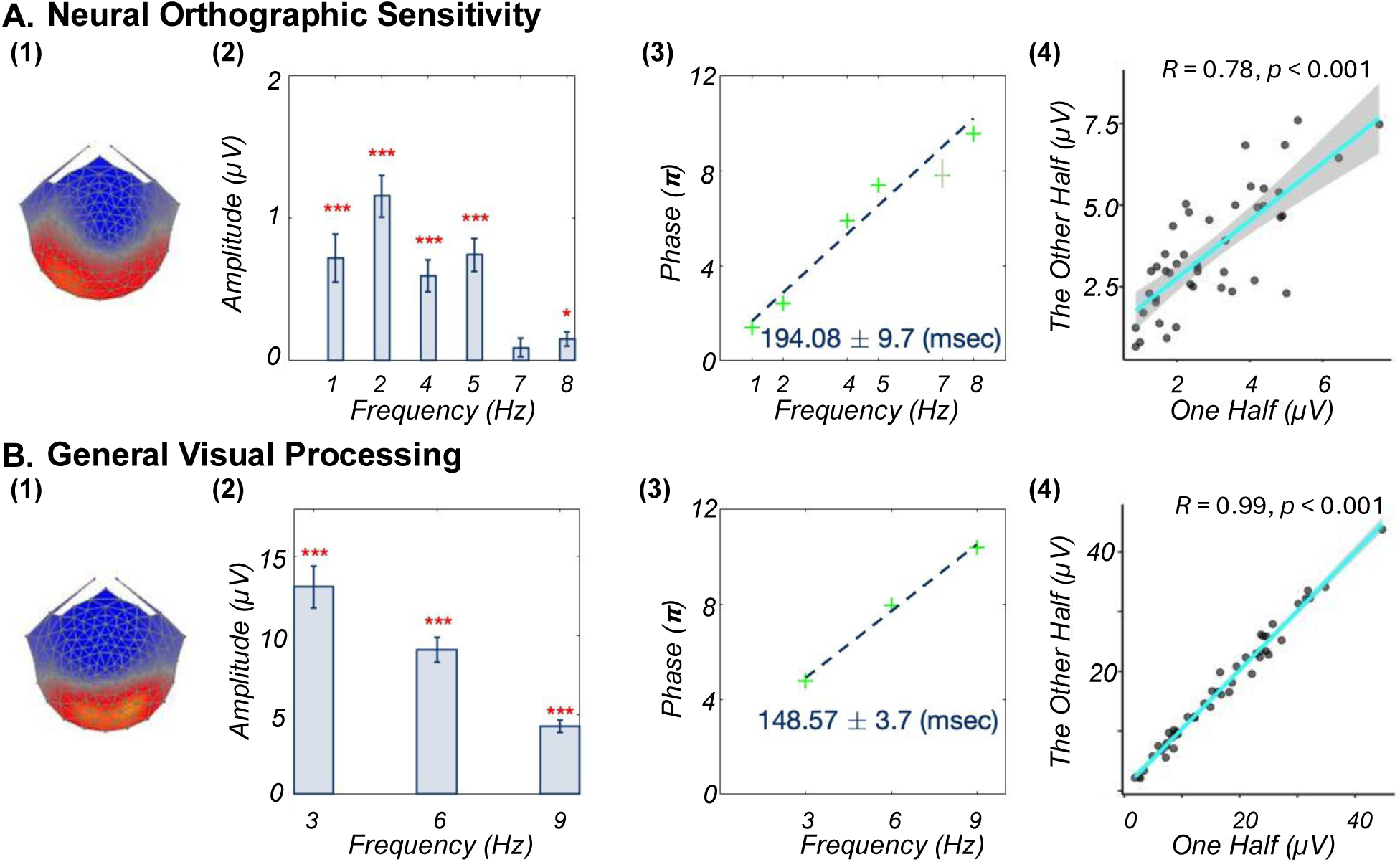
Robust neural responses to onset–rime structure and general visual information. A: Oddball responses reflecting neural sensitivity to onset–rime structure. (1) Topographic maps and (2) corresponding bar plots show response amplitudes across harmonics over left occipitotemporal regions; (3) Phase (*π*) by harmonic frequency functions were used to estimate response latency in milliseconds (ms), yielding a latency estimate of 194.08 *±* 9.7 ms; (4) Split-half reliability analysis demonstrating the stability of orthographic sensitivity responses and ruling out potential confounds arising from different stimulus sets. B: Base responses reflecting general visual processing. (1) Topographic maps and (2) corresponding bar plots show response amplitudes across harmonics over occipital visual regions. (3) Phase (*π*) by harmonic frequency functions yielded a latency estimate of 148.57 *±* 3.7 ms. (4) Split-half reliability analysis demonstrating the stability of base visual responses. ***: *p_F DR_ <* 0.001, *: *p_F DR_ <* 0.05.

Figure 2B demonstrates neural responses to general visual processing. Topographies (Figure 2B.1) and bar plots (Figure 2B.2) illustrate significant neural responses over occipital visual regions at all base harmonics. Phase (*π*) by harmonic frequency functions yielded an estimated latency of 148.57 *±* 3.7 ms (Figure 2B.3), which seems more consistent with a P1-like visual response in the ERP field. Figure 2B.4 demonstrates high split-half reliability of the general visual response.

### 3.2 Neural sensitivity to onset–rime structure, but not general visual processing, is associated with spelling and reading

Oddball RSS amplitude was significantly associated with both WJ Spelling (*r* = 0.62*, p_F DR_ <* 0.001, Figure 3A.1) and TOWRE (*r* = 0.53*, p_F DR_ <* 0.001, Figure 3A.2), indicating that greater neural sensitivity to onset–rime structure was associated with better spelling ability and word-reading efficiency. These associations remained significant after removal of influential observations identified by Cook’s Distance. Multiple linear regression analyses further showed that the associations between RSS amplitude and WJ Spelling (*β* = 0.48*, t* = 3.13*, p* = 0.003) and TOWRE (*β* = 0.54*, t* = 3.77*, p <* 0.001) remained significant after controlling for age.

**Figure 3:**
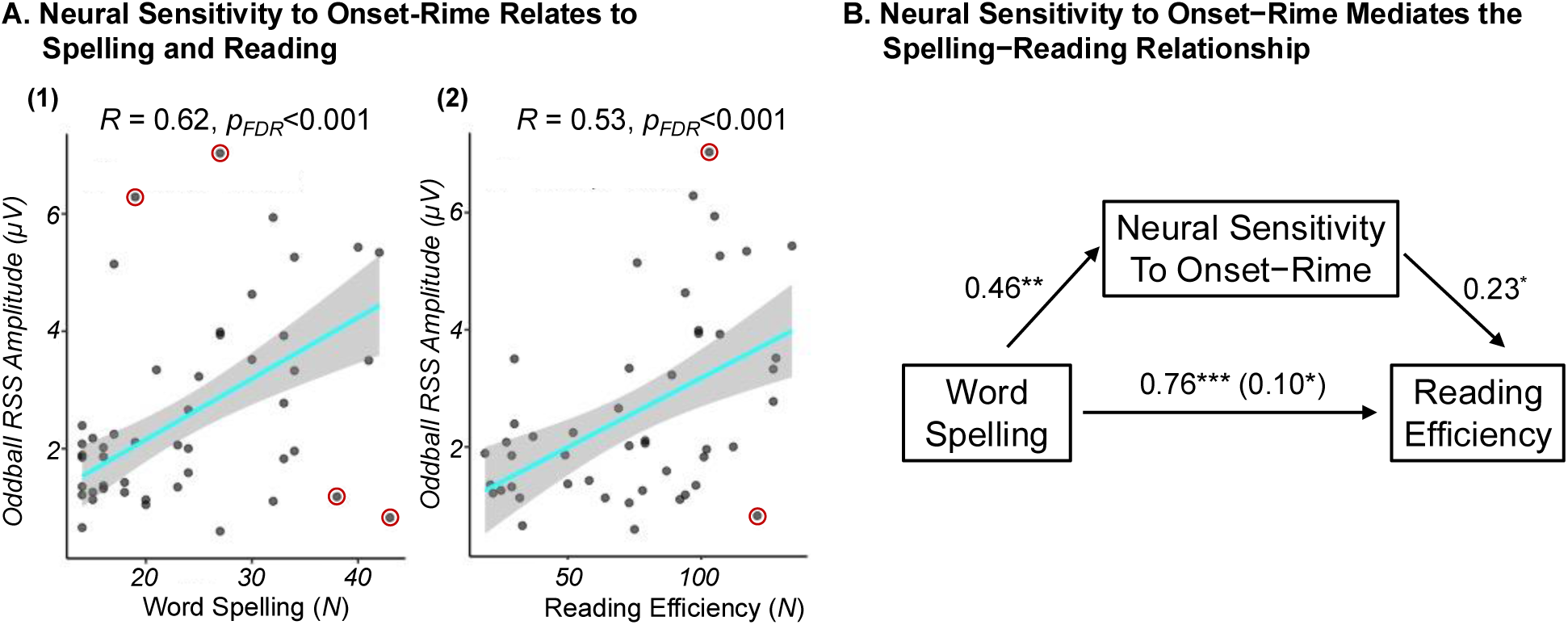
Neural sensitivity to onset–rime structure is associated with reading and spelling and mediates the relationship between these two literacy skills. A: Linear regressions between Oddball RSS amplitude across all harmonics and spelling ability (measured by WJ Spelling) as well as word-reading efficiency (measured by TOWRE). Children with greater neural sensitivity to onset–rime structure showed better spelling ability and word-reading efficiency. These relationships remained significant after controlling for age and after removing influential observations (circled in red). B: Mediation analyses revealed a significant indirect pathway linking spelling ability and word-reading efficiency through neural sensitivity to onset–rime structure.. Specifically, better spelling ability was associated with greater neural sensitivity to onset–rime structure, which in turn was associated with greater word-reading efficiency. WJ: Woodcock- Johnson; TOWRE: Test of Word Reading Efficiency.***: *p_F DR_ <* 0.001, **: *p_F DR_ <* 0.01,*: *p_F DR_ <* 0.05.

To assess whether these brain–behavior relationships were specific to neural sensitivity to onset–rime structure, we additionally examined correlations between RSS amplitude of the base responses and both WJ Spelling and TOWRE scores. In contrast to the oddball response, no significant correlations were found between base RSS amplitude and either behavioral measure (all *r <* 0.24*, p >* 0.11). These findings indicate that the observed brain–behavior relationships were specific to neural sensitivity to onset–rime structure rather than general visual processing.

### 3.3 Neural sensitivity to onset–rime structure mediates the association between spelling and reading

Given the observed associations between neural sensitivity to onset–rime structure and both behavioral measures, mediation analyses were conducted to test whether this neural sensitivity mediated the relationship between spelling ability and word-reading efficiency. Results (Figure 3.B)indicated a significant indirect effect of spelling ability on word-reading efficiency through neural sensitivity to onset–rime structure. The standardized regression coefficient between word spelling and reading efficiency (*β* = 0.76*, p_F DR_ <* 0.001), as well as the standardized regression coefficient between neural orthographic sensitivity and reading efficiency (*β* = 0.51*, p_F DR_ <* 0.001), were both significant. The standardized indirect effect was (0.46)×(0.23) = 0.10. We tested the significance of this indirect effect using bootstrapping procedures (1000 samples). The bootstrapped indirect effect was 0.10, with a 95 % confidence interval of [0.02, 0.19], and was statistically significant (*p_F DR_ <* 0.05). The total effect of spelling ability on word-reading efficiency was also significant (*β* = 0.76*, p_F DR_ <* 0.001), indicating that word spelling influences reading efficiency both directly and indirectly through neural orthographic sensitivity.

We further examined whether neural sensitivity to onset–rime structure explained additional variance in word-reading efficiency beyond spelling ability and age. Hierarchical regression analyses showed that adding oddball RSS amplitude to a model including WJ Spelling and age resulted in a significant increase in explained variance in TOWRE performance (Δ*R*^2^ = 0.04*, F* (1, 43) = 8.29*, p* = 0.006). Thus, neural sensitivity to onset–rime structure accounted for unique variance in word-reading efficiency beyond spelling ability and age.

## 4 Discussion

### 4.1 Neural Evidence for Onset–Rime Structure as an Intermediate Orthographic Representation

In first- and second-grade children, neural sensitivity to onset–rime structure was characterized by a component distributed over the left occipito-temporal (OT) region. This aligns with previous ERP, SSVEP, and fMRI studies demonstrating neural sensitivity to orthographic information over the left OT, a region known to play a critical role in fluent and efficient reading (Dehaene & Cohen, 2011; Lochy et al., 2016; McCandliss et al., 2003; Wang et al., 2023; Zhao et al., 2014). For example, Zhao et al. (2014) found ERP response differences between orthographically legal pseudowords and orthographically illegal consonant strings at left OT electrodes in 7-year-old children.

Most studies to date have focused on neural responses to word form (words vs. pseudo-fonts), lexicality (words vs. pseudowords), and simple orthographic regularities (e.g., orthographically legal vs. illegal letter strings), leaving open the question of whether early readers encode intermediate orthographic representations. For instance, a recent SSVEP study (Wang et al., 2023) reported robust neural sensitivity to orthographic structure in early readers using consonant–vowel–consonant (CVC) letter strings (e.g., yot vs. igd). However, this manipulation tested the legality of local letter sequences rather than the organization of larger recurring orthographic patterns. In contrast, behavioral studies have shown that developing readers are sensitive to larger orthographic structures such as onset–rime organization (Goswami, 1991; Goswami & Mead, 1992; Greaney et al., 1997; Wise et al., 1990). Yet little is known about how such intermediate orthographic structure is represented in the developing brain.

The present findings extend this behavioral literature by providing neural evidence that children in the early stages of literacy development are sensitive to onset–rime organization even when no explicit judgment of orthographic structure is required. The observed neural discrimination between onset–rime and rime–onset strings suggests that this organization is encoded within children’s visual orthographic representations, consistent with the proposal that onset–rime structure may form an intermediate representation between local letter combinations and whole-word forms. Our previous FPVS work showed that early readers are neurally sensitive to orthographic structure in short CVC strings (Wang et al., 2023), and subsequent longitudinal work showed that individual differences in this neural sensitivity were associated with reading skill and that its development over time was related to growth in reading fluency (Wang et al., 2024). The present study builds on these findings by showing that neural orthographic sensitivity extends beyond local letter-sequence regularities to the organization of larger, reusable orthographic units. This provides a more specific account of what we previously characterized broadly as visual orthographic sensitivity: part of this sensitivity may reflect how developing readers represent recurring orthographic structures that are shared across words.

Importantly, the observed neural responses to onset–rime structure were highly reliable, as demonstrated by strong split-half reliability across different blocks of stimulus sets. This finding suggests that the measured responses reflect a stable sensitivity to onset–rime organization rather than being driven by the specific items included in a given stimulus set. Moreover, sensitivity to onset–rime structure cannot be explained simply by familiarity with particular visual word forms. The onset–rime and rime–onset strings were constructed from the same orthographic elements but differed in how those elements were ordered. The neural discrimination therefore cannot be attributed solely to exposure to the individual elements themselves, but instead depends on how those elements are organized.

Phase-lag quantification of the component yields an estimated latency of approximately 190 ms. This 190 ms latency is consistent with a previous ERP study in 7-year-old children reporting sensitivity to orthographic properties at around 200ms (Zhao et al., 2014), slightly later than that in adults at around 150–180 ms (Araújo et al., 2015). The slightly shorter latency in adults might reflect increased efficiency of orthographic processing with reading experience and age (Maurer et al., 2005). This interpretation is consistent with recent FPVS evidence showing that children with shorter cortical latencies tend to exhibit better reading fluency and that latency decreases with age across late childhood and early adolescence (Wang et al., 2025).

In addition to the neural responses observed for onset–rime oddballs, we also identified a robust RCA component that was maximal over bilateral occipital regions during rime–onset base stimulation. Base-frequency responses in FPVS paradigms are generally thought to reflect visual processing of the stimulus stream itself (Lochy et al., 2016), whereas oddball responses index sensitivity to differences between the base stream and periodically presented oddballs (here, onset–rime vs. rime–onset). The bilateral occipital activation observed here is consistent with previous FPVS studies especially those using similar stimulation rates (Wang et al., 2023, 2025). This scalp topography also corresponds to expected activations reported in fMRI literature (López-Barroso et al., 2020; Turkeltaub et al., 2003b; Szwed et al., 2011) and in EEG source localization studies (Rossion et al., 2003; Proverbio & Adorni, 2009). The estimated latency of approximately 150 ms is also consistent with early visual processing typically observed within this time range (Brem et al., 2006; Maurer et al., 2006; Proverbio & Adorni, 2009). Similar to the onset–rime oddball response, the base response showed exceptionally high split-half reliability (*R* = 0.99), further demonstrating the reliability of the FPVS paradigm in capturing both general visual processing and sensitivity to orthographic structure.

### 4.2 Neural Sensitivity to Onset–Rime Structure, but Not General Visual Processing, Is Associated With Reading and Spelling

Among beginning readers, neural sensitivity to onset–rime orthographic structure was positively associated with both spelling and reading. Children with stronger neural orthographic sensitivity had better spelling and reading skills, even after controlling for age and removing influential outliers. These findings suggest that individual differences in sensitivity to this intermediate orthographic representation are relevant to both reading and spelling skills.

These results are consistent with previous behavioral findings linking orthographic processing to reading and spelling development. For example, Rakhlin et al. (2019) identified orthographic processing as a key predictor of reading fluency. Likewise, orthographic knowledge has been shown to contribute to spelling accuracy and spelling development beyond phonological skills (Conrad et al., 2019; Treiman, 2017). Similar associations have also been observed at the neural level. FPVS studies have shown that neural sensitivity to orthographic structure is positively correlated with reading skill in typically developing readers (Wang et al., 2024), while reduced orthographic discrimination responses have been observed in children with reading difficulties (Lutz et al., 2024).

Sensitivity to recurring orthographic patterns, such as onset–rime structure, may be particularly important during early reading acquisition in English, where phoneme–grapheme correspondences are often inconsistent. Larger recurring patterns can provide more consistent mappings across orthographically related words and may therefore offer a useful basis for learning and applying familiar spelling patterns (Goswami, 1991; Ehri, 2000).

One possible explanation for the observed associations is that neural sensitivity to on-set–rime structure reflects the development of more refined visual orthographic representations. As children gain reading experience, they gradually learn that written words are not simply collections of individual letters but contain recurring patterns that can be shared and generalized across words (Deacon et al., 2019; Ehri, 2014; Treiman, 2020). Sensitivity to onset–rime structure may therefore support more efficient recognition of familiar words, decoding of unfamiliar words, and generalization of orthographic knowledge across related words.

The fact that only neural sensitivity to onset–rime structure, but not general visual processing of the stimulus stream, was correlated with reading and spelling further suggests that literacy skills are associated with children’s sensitivity to meaningful orthographic organization rather than general visual processing of written stimuli.

### 4.3 Neural Orthographic Sensitivity Mediates the Association Between Spelling and Reading

A central finding of the present study is that neural sensitivity to onset–rime orthographic structure mediated the association between spelling and reading. Whereas the close relationship between these two literacy skills has been well established behaviorally (Ehri, 2000; Georgiou et al., 2020), the neural representations that may contribute to this relationship remain poorly understood. Specifically, stronger spelling ability was associated with greater neural sensitivity to onset–rime structure, which in turn was associated with more efficient word reading.

The observed mediation may help explain how orthographic knowledge contributes to the relationship between spelling and reading. Spelling requires learners to generate complete written forms by accurately encoding and retrieving the identities and positions of letters within a word (Perfetti, 1997; Treiman et al., 2019), which may encourage children to attend to the detailed orthographic structure of written words (Conrad et al., 2019; Ouellette, 2010). Spelling may therefore place greater demands on orthographic knowledge than reading (Treiman, 2017; Conrad et al., 2019; Ouellette, 2010). In line with this, longitudinal and intervention studies further suggest that spelling activities facilitate orthographic learning, which subsequently supports reading development (Conrad et al., 2019; Ouellette & Śeńechal, 2017). Through repeated spelling experience, children may become increasingly sensitive not only to individual letters and letter combinations but also to how these elements are organized into larger orthographic structures, such as onset and rime units, that recur across words (Conrad et al., 2019; Goswami, 1991). The present findings suggest that sensitivity to onset–rime structure may be an important component of orthographic knowledge during the early stages of literacy acquisition. Such sensitivity may, in turn, support reading by facilitating the recognition and decoding of familiar words and the application of orthographic knowledge to unfamiliar but related words.

The present findings also extend previous behavioral accounts proposing that the close relationship between spelling and reading arises, in part, from their reliance on shared orthographic knowledge (Chang et al., 2018; Mather & Jaffe, 2021). However, behavioral measures based on overt decisions and responses provide limited insight into the nature of visual orthographic representations underlying this relationship. Here, neural sensitivity to onset–rime structure was measured while the children performed an unrelated sea-animal detection task, without requiring explicit judgments about orthographic structure. Moreover, robust neural discrimination was measured from brief periods of stimulation, providing an objective index of rapid and automatic sensitivity to orthographic organization rather than children’s explicit ability to report or manipulate that knowledge. Although the present findings do not establish that reading and spelling recruit *identical* neural representations, the mediation effect is consistent with the possibility that intermediate orthographic representations provide a neural link between the two skills.

### 4.4 Limitations

Several limitations of the present study should be noted. First, we designed the onset–rime vs. rime–onset contrast to isolate sensitivity to onset–rime organization; however, this manipulation may also capture other orthographic properties that coincide with onset–rime organization. For example, onset–rime strings are generally more word-like and pronounceable than their rime–onset counterparts and may better reflect the way these orthographic elements are normally arranged in English words. As a consequence, the present design cannot fully determine whether the observed responses reflect sensitivity specifically to on- set–rime structure or to broader orthographic properties associated with this organization. Nevertheless, because the two stimulus types were composed of the same onset and rime elements and differed only in their arrangement, the findings provide evidence that children are sensitive to how larger orthographic units are organized within written words. Future work will be needed to better characterize the specific orthographic properties that contribute to this sensitivity. Second, although the mediation analysis suggests that neural sensitivity to onset–rime structure may help explain the relationship between spelling and reading, the cross-sectional design does not allow strong conclusions about causal or developmental direction. Longitudinal studies are needed to determine whether spelling experience contributes to the development of neural sensitivity to onset–rime structure and whether such sensitivity subsequently supports reading growth. Finally, future studies with larger and more diverse samples will be important for testing the robustness and generalizability of these findings.

Despite these limitations, the present findings provide neural evidence that developing readers are sensitive to onset–rime organization and suggest that this sensitivity may reflect an intermediate orthographic representation between individual letters and whole-word forms. Moreover, neural sensitivity to onset–rime structure was associated with both reading and spelling and mediated the relationship between these two literacy skills, providing further evidence that intermediate orthographic representations may contribute to their close relationship.

## 5 Conclusion

The present findings provide neural evidence that first- and second-grade children are sensitive to intermediate orthographic structures that extend beyond individual letters and smaller letter combinations but fall below the whole-word level. Building on behavioral evidence for the role of onset–rime structure in reading and spelling, the present study demonstrates robust and automatic neural sensitivity to this structure during early literacy development. Critically, neural sensitivity to onset–rime structure mediated the relationship between spelling and reading. Together, these findings provide a possible neural account of why spelling and reading are so closely related: the two skills may become linked through intermediate orthographic representations that capture how smaller orthographic elements are organized into larger units that recur across words. This possibility may also have implications for instruction, suggesting that attention to how letters and letter combinations form recurring larger units could support orthographic knowledge that is useful for both spelling and reading.

